# LoMuS: Low-Rank Adaptation with Multimodal Representations Improves Protein Stability Prediction

**DOI:** 10.64898/2025.12.15.694540

**Authors:** Samuel Infante, Akash Singh, Anowarul Kabir

## Abstract

Protein folding stability is a key determinant for understanding protein dynamics, including molecular function, pathogenicity, and/or protein engineering. Yet, accurate prediction of protein stability changes remains a challenging problem due to the high-variability in the available data, especially from sequence-only information when structural knowledge is of low-resolution or unavailable. In this work, we introduce LoMuS, a <u>Mu</u>ltimodal deep learning model that combines two complimentary aspects of the molecule and predicts unnormalized protein <u>S</u>tability effect from the primary sequence as input. In the core of the model architecture, a fusion network integrates explicit physicochemical descriptors with <u>Lo</u>w-rank adapted protein language model derived embeddings from the sequence that shows powerful and accurate generalization ability across various benchmark settings for predicting protein folding stability changes. We compared and rigorously evaluated our model capacity spanning from fold-induced stability changes to mutation caused stability effect prediction. This includes benchmarking against various held-out protein domains, out-of-distribution label settings and per-protein evaluation. LoMuS consistently outperforms other sequence-only protein stability baselines. It achieves an absolute performance gain by an at least 10% in the spearman rank correlation metric for predicting protein stability across many held-out domains and out-of-distribution stability label predictions. Per-protein validation additionally demonstrates promising performance gain of our model. Ablation analysis on the model architectural choices confirms that complementary signals from derived features are critical for this multimodal approach. We believe LoMuS advances protein engineering research and can aid in rational protein design by elucidating precise protein stability changes.

**Availability:** All codes including data preparation scripts, training and validation recipes, and experimental configurations for LoMuS are available at: https://github.com/samuelinfantee/LoMuS-repository.

**Supplementary information:** Supplementary data are available at *Journal Name* online.

## Introduction

Reliable protein stability prediction has the potential to transform biotechnology, human health, drug repurposing and medical therapeutics. Stable proteins are often used as enzymes in biopharmaceuticals and safe industrial food [1]. In contrast, unstable proteins, caused by missense mutations, can lead to many human diseases such as Alzheimer, cystic fibrosis, and Lynch syndrome [2, 3, 4], since many pathogenic variants act by lowering protein stability [5]. By elucidating the core in the sequence-structure-function relationship, accurate prediction of protein folding stability can aid in rational design in protein engineering and variant interpretation [6].

Protein stability generally refers to the free-energy gap between native (*G*_*N*_ ) and unfolded (*G*_*U*_ ) ensembles under certain conditions, defined as Δ*G*_*fold*_ = *G*_*N*_ − *G*_*U*_. This folding stability can also be changed due to the mutations, measured as 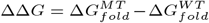 where 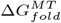 and 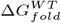 indicates free-energy gap for mutant- and wild-type variants of the same protein molecule, respectively. Here, depending on the ΔΔ*G* equation, positive and negative values denote destabilization and stabilization, respectively. Experimental Δ*G* and ΔΔ*G* are typically derived under two-state assumptions from chemical denaturation or thermal unfolding and mapped to free-energy with standard analyses [7, 8]. Being low-throughput, laborious, and expensive, such methods are not widely applicable across many proteins or large-scale variants screening. To address this problem, several computational methods have been proposed over the years; however, they often suffer from scaling and generalizability due to the out-of-distribution problem [9] often raised in the biological domain.

Classical methods are computationally expensive and limited in scale due to the usage of empirical energy functions based on physical chemistry, such as FoldX [10] and Rosetta [11]. They compute ΔΔ*G* from changes in molecular energy using force fields and free-energy centric simulations, makes them unamenable even for a modest number of variants [12]. Additionally, methods relying on protein three-dimensional structures are not applicable when such structures are not available [13]. Consequently, classical models struggle with large-scale mutational scanning and often not usable for long proteins.

Machine learning (ML) approaches train statistical methods on existing stability data. mCSM [14] models the residue environment using graph-based signals to predict stability changes. DeepDDG [15], an early neural-network model, uses mutation centric residue features trained on *∼* 5, 700 mutations to predict mutation-induced stability. Ensemble methods like MAESTRO [13] combine multiple regression agents, and newer tools like PremPS [16] employ random forests with structure- and evolution-based features. Most ML models rely on limited curated ΔΔ*G* datasets, and high-quality structures for feature extraction. They often tend to overfit common protein families and offer only modest accuracy for novel cases.

DeepDDG, for instance, achieved pearson’s correlations of only *∼* 0.5–0.6 on independent test sets [15]. Another issue is that most predictors systematically compress their ΔΔ*G* outputs toward zero, which means that they underestimate large effects [17]. In particular, stabilizing mutations (large negative ΔΔ*G*) are often missed [17]. PremPS explicitly balanced its training set to improve stabilizing predictions; however, it still reports degraded accuracy on low-resolution or homology-model structures [16]. Moreover, Laimer et al. [13] observed that the scarcity and bias of training data (many mutations from few proteins) can cause ML predictors to overfit; for proteins absent from the training set, performance drops notably.

Although data-driven methods have improved folding stability prediction, recent deep learning-based advancements still inherit key constraints. CNNs on structural micro-environments (RaSP [18]) and graph-based models that fuse sequence embeddings with 3D information (ELASPIC-2 [19]) report improved performance. PMSPcnn [20] uses persistent homology extracted topological features to predict stability effect caused by point mutations and improves over prior baselines. Hybrid designs continue this trend, for example DDMut [21], a siamese network that integrates graph encodings of local 3D context with convolutional and transformer layers, and ProS-GNN [22] uses message passing on atomistic graphs to capture both short- and long-range effects. All these methods rely on structural information and curated labels, which makes them limited in application when no structures are available or structures are of low resolution. However, the cumulative strengths of these works suggest that a more suitable method focusing on sequence representations with lightweight but broadly available biochemical signals might be of great interest when structural data are scarce.

Recent trend leverages protein language models (PLMs) pretrained on a large corpus of protein sequences [23, 24, 25]. They provide powerful general-purpose embeddings that can be transferred into diverse relevant prediction tasks, including secondary structure, disorder, solvent accessibility, mutation effect, function and stability [26, 27, 28]. Crucially, Schmirler et al. [29] showed that supervised fine-tuning of PLMs almost always improves downstream protein prediction tasks, however might subject to overfitting [30]. Transformer PLMs can capture implicit structural information and enable accurate tertiary structure prediction from a single sequence without MSAs [31]. Another method, augmented with structural or physicochemical inputs combined with PLM embeddings, reported improved performance for protein stability change (ΔΔ*G*) estimation [32]. Beyond discriminative settings, generative and task-conditioned LLMs have demonstrated practical impact in engineering workflows, for example a temperature-guided model proposing stabilizing mutations with more than 30 percent experimental success across multiple proteins [33]. Another class of parameter-efficient methods such as low-rank adapters (LoRA) can achieve on-par performance while cutting trainable parameters [34]. Overall, these results suggest that careful task-specific adaptation can lead to capturing subtle stability changes caused by mutation.

Considering the drawbacks and progress discussed earlier, we propose a deep learning based computational approach that considers only protein sequence as input, no dependency on structures or multiple sequence alignments, and leverages a multimodal architecture that makes the model more-generalizable across different benchmarks. Here, we introduce LoMuS, a <u>Lo</u>w-rank adaptation of embeddings with <u>Mu</u>ltimodal protein representation to predict protein <u>S</u>tability changes, for both Δ*G* and ΔΔ*G* based on the available datasets. LoMuS consists of three core components, (i) global physicochemical features that we derive from the given sequence, (ii) a LoRA adapted PLM that we finetune in-house by freezing most weights and only learning small rank-decomposition adapters in each Transformer layer [34], and (iii) a fusion network where PLM embeddings and the global representation are fused and passed through a lightweight neural head. Intuitively, the model learns to weight the contribution of the learned sequence embedding versus the physicochemical features.

We train LoMuS on experimental datasets and evaluate its performance across three settings: (i) generalization across held-out domains, (ii) generalization across out-of-distribution stability labels, and (iii) held-out proteins. The resulting model leverages the complementary strengths of sequence-based and descriptor-based learning, achieving state-of-the-art accuracy on several stability benchmarks. LoMuS outperformed its closest baseline by an absolute *∼* 10% in the spearman’s rank correlation metric for predicting protein stability across many held-out domains. In another direction, for out-of-distribution stability label generalization task, our approach consistantly achiches higher performance by an absolute *∼* 13%. On per-protein mutation caused stability prediction generalization task, LoMuS matches or exceeds a strong sequence-only reference. Taken together, these results show that fusing a LoRA-adapted PLM with compact physicochemical descriptors improves ranking accuracy for stability both within families and across held-out domains. In summary, LoMuS combines rich PLM embeddings with simple biophysical inputs (a strategy that captures both learned sequence context and known protein chemistry) to improve mutation stability predictions.

In the rest of the article, we discuss our methodological contributions and subsequently we report and compare our results with the state-of-the-art methods. Section 2 presents LoMuS, including the LoRA fine tuning strategy, physicochemical feature set, fusion network, and training setup. We then summarize the datasets and preprocessing steps in Section 3. Section 4 reports results with standard performance metrics and ablation studies to understand various model choices. Finally, in Section 5, we outline the drawbacks that we deem as the future directions for extending this line of research.

## Methodology

### LoMuS Overview

We propose a multimodal deep learning architecture, namely LoMuS, that estimates the stability score of a protein molecule from the primary sequence as input. Our model integrates three sets of features: (1) sequence embeddings generated by a pretrained PLM, (2) physicochemical descriptors and (3) basis features, summarized in Fig. 1. We leverage the ESM-2 transformer-based encoder [23] and fine-tune it using Low-Rank Adaptation (LoRA) [34] while preserving most pretrained weights to extract sequence embeddings. The physicochemical features are derived from AAindex [35, 36, 37, 38]. Subsequently, we compute six basis descriptors of the input molecule. Finally, our designed fusion network combines all of those features and accurately predict the stability scores. The following subsections describe each component of our architecture in detail.

**Fig. 1.**
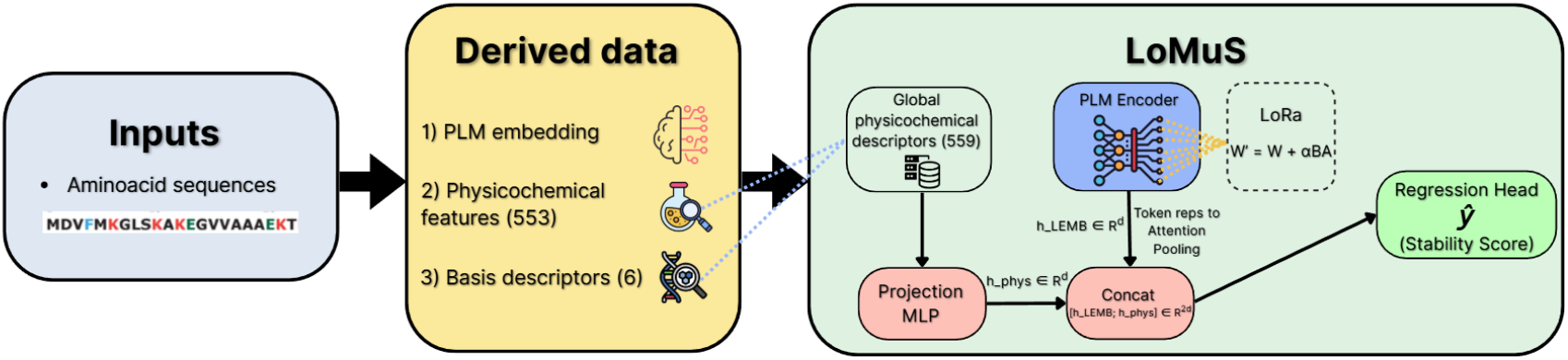
Procedural overview of LoMuS architecture. Given amino acid sequence as input, LoMuS derives three sets of features: (i) embeddings generated from a pretrained protein language model, (ii) physicochemical descriptors, and (iii) basis features, including length, molecular weight, aliphatic index, instability index and so on. The sequence is encoded by a pretrained PLM and adapted with LoRA on the attention projections. In parallel, physicochemical feature vector is standardized and mapped by a multi-layer perceptrons (MLPs) into the same PLM feature dimension. The model concatenates both and feeds it to a lightweight regression head that outputs a stability score. The target is Δ*G* or ΔΔ*G* depending on the dataset. This fusion lets the network weight information from the learned sequence embedding and the explicit physicochemical context during end-to-end training.

### From Input Sequence to Learned representation

We use the ESM-2 [23] (650M parameters) to extract the learned representation of the input protein sequence. ESM-2 is pretrained on the UniRef50 [39] corpus comprising billions of protein sequences. With multiple self-attention layers, it is designed to capture implicit patterns in sequences correlated with underlying structure and function. Each sequence is then fed into the ESM-2 encoder to produce a contextual embedding for each residue. Given an input amino acid sequence of length *L*, ESM-2 (hidden size *d* = 1280) yields embeddings *X* ∈ ℝ^*L×d*^.

### Learning Task-relevant Features using Low-Rank Adaptation (LoRA) Fine-Tuning

To adapt the PLM to our task-of-interest, stability prediction, we apply Low-Rank Adaptation (LoRA) [34] to the model’s attention layers. LoRA is a parameter-efficient fine-tuning technique (PEFT) that freezes the original pretrained weights and injects trainable low-dimensional matrices into each layer’s weights. This reduces the overfitting risk under limited labeled data and retains PLM knowledge for improving out-of-distribution generalization performance [40], as in ours.

In practice, for each attention projection weight matrix *W* in the PLM (query *Q*, key *K*, value *V*, and output), we introduce two low-rank matrices *A* ∈ ℝ^*d×r*^ and *B* ∈ ℝ^*r×d*^ with *r ≪ d*. The adapted weight is formulated in Eqn. 1.

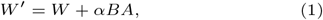

where *α* is a scaling factor that controls the initial magnitude of the LoRA update. Here, *W* remains fixed, and only *A* and *B* are learned, greatly reducing the number of tunable parameters [34] in our model. We apply LoRA to the self-attention projection matrices *Q, K, V*, and the output projection in all encoder layers following [41], while leaving the feed-forward blocks and the embedding layer frozen. This PEFT approach retains ESM-2’s implicitly learned evolutionary and structural features while allowing LoMuS to adjust the attention dynamics for stability prediction with minimal overhead. We set the value LoRA rank *r* and scaling *α* based on preliminary experiments in [42, 43], specifically *r* = 9, *α* = 16. During fine-tuning, only the LoRA parameters and subsequent new layers are updated; the vast majority of PLM’s original parameters remain fixed, avoiding overfitting and reducing GPU memory requirements.

### Multimodal Representation of a Protein Molecule: Physicochemical and Basis Features

LoMuS incorporates local physicochemical properties and global sequence-level descriptors in addition to sequence embeddings. We derive these features from each sequence *a priori*. The AAindex1 database is used to extract the physicochemical properties of each amino acid in the sequence [35]. This feature set consists of 553 published indices, where each index provides a numeric value for each 20 canonical amino acid, representing physicochemical properties, such as hydrophobicity, flexibility, polarizability, chemical shifts and many. We apply an average pooling across the sequence to compute physicochemical representation of the molecule, yielding a 553-dimensional vector for the sequence. These features summarize the overall tendency of the sequence with respect to a wide range of biochemical scales.

Next, we derive six global physicochemical descriptors frequently used in protein property and stability modeling [44, 45] of a protein molecule. These include sequence length, molecular (MW) weight, isoelectric point (IP), grand average of hydropathy (GRAVY), aliphatic index (AI) and instability index (II) (More in Supplementary S1.2). These six features are derived directly from the raw amino acid sequence using Biopython’s ProteinAnalysis toolkit, implemented in line with ExPASy ProtParam conventions [46, 47]. Finally, we have the physicochemical representation of the protein molecule by concatenating all features as in Eqn. 2.

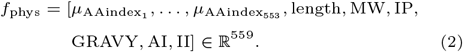

### LoMuS Fusion Architecture

After extracting the LoRA enabled residue contextual embeddings from the PLM and the global physicochemical feature vector, LoMuS employs a fusion network to predict the protein stability score. The core components of the network are discussed in the following subsections.

#### Protein representation generation from amino acid level embeddings

We apply an attention pooling operation over the residues to compute the protein level representation. Let *H* = (*h*_1_, …, *h*_*L*_) denotes the residue embeddings extracted from the last PLM layer, where *L* is the input sequence length. The relevance score for residue *i* is determined by a learnable linear projection (*W*_attn_) layer followed by a tanh nonlinearity, and a learned context vector *u* as in Eqn. 3:

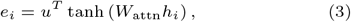

Subsequently, attention weights (*α*_*i*_) for each residue *i* are obtained by excluding padded positions by masking with a very large negative number, then normalizing the scores across positions with a softmax function, in Eqn. 4:

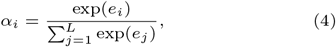

The sequence-level representation is a weighted average of the residue embeddings. If the weights concentrate on a few positions, the representation emphasizes those key residues.

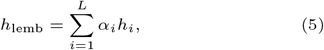

The physicochemical vector *f*_phys_ ∈ ℝ^559^ is mapped into the same latent space as the PLM embedding dimension via a multi-layer perceptron *g*_*w*_(·) with nonlinearity ReLU activation function and dropout (rate=0.1):

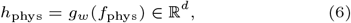

where *d* is the PLM hidden size (i.e., *d* = 1280 for ESM-2). *g*_*w*_(·) aligns scale and geometry of physicochemical features within the PLM space, stabilizing fusion and allowing richer interactions in the downstream tasks.

#### Fusion of Multimodal Features

We concatenate the sequence level LoRA enabled PLM embedding (*h*_lemb_) and projected physicochemical features (*h*_phys_) to form combined embedding as in Eqn. 7.

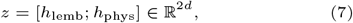

where [*x*; *y*] denotes vector concatenation. The fused vector *z* captures both sequence-derived patterns from the LoRA-PLM and global physicochemical properties.

#### Loss/objective Function

The model deploys a regression head to estimate the stability score *ŷ* given the fused sequence-level vector representation *z*. We train the model end-to-end by minimizing the mean-squared error (MSE) between predictions *ŷ* and ground truth labels *y*. For a mini-batch of *n* samples, the loss can be defined as:

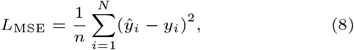

which provides a smooth regression objective to drive learning. In Fig. 2, we report the validation rank correlation (spearman’s *ρ*) as a function of training epoch across various benchmark datasets, discussed in the Datasets 3 section. All benchmarks show similar trend of rapid gains in the first few epochs followed by a shallow plateau, no overfitting, indicating early convergence under a fixed training configuration.

**Fig. 2.**
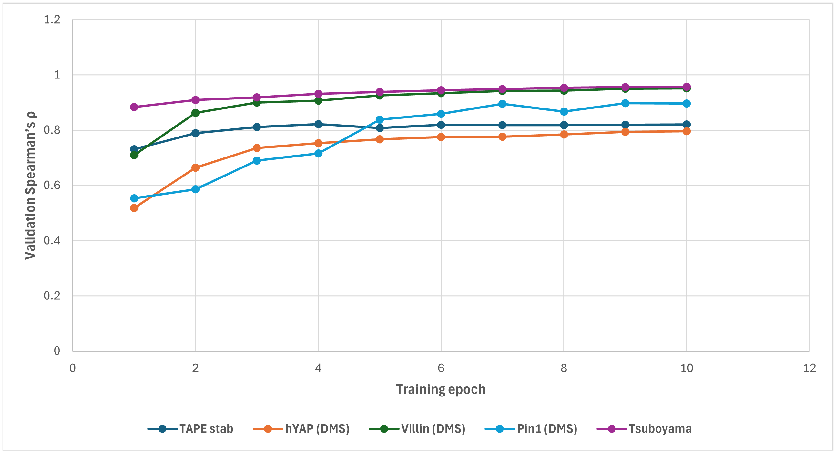
Evolution of validation Spearman’s *ρ* across training epochs on various benchmark datasets.

### Training Details

We train LoMuS end-to-end using the AdamW optimizer, keeping the pretrained backbone weights fixed and updating only the LoRA adapters. To account for the differing roles and sizes of the components, we use two learning rates: a higher rate for the PLM branch, namely the LoRA-adapted attention projections, and a lower rate for the newly initialized fusion and regressor layers. This setup encourages the backbone’s attention projections to adapt meaningfully to the task while keeping the smaller fusion head conservative to avoid overfitting to the handcrafted features. In our experiment, we found that setting the PLM+LoRA learning rate larger (for example, 5*e* − 4) than the fusion and regressor learning rate (for example, 1*e* − 4) works well.

We also employ a short linear warm-up schedule at the start of training, where the learning rates ramp from 0 to their targets over the first few epochs or early percentage of steps, to stabilize optimization under the mixed frozen and adapter regime, and we apply dropout at 0.1 in the projection and pooling components. After warm-up, we either hold the rates constant or decay them slightly based on validation performance.

To resolve overfitting, we apply regularization during the training. We apply gradient clipping (clamping the gradient norm to a fixed threshold) to avoid any outlier updates that could destabilize the LoRA weights or the MLPs. We train for a fixed number of epochs, evaluating on a validation set at regular intervals. Model checkpoints are saved, and early stopping is triggered based on validation set performance (spearman’s rank correlation). This selection criterion aligns with the goal of getting the relative stability ranking correct, which is often important in protein engineering. MSE on the validation set is also monitored to ensure the model is not overfitting.

Once training is complete, the final model, which combines the LoRA-adapted PLM weights with the trained MLPs, is used to predict stability on test proteins. The physicochemical feature scaler fitted on the training data is applied to normalize new sequences’ features, and tokenization is done with the same PLM vocabulary to ensure consistency. The result is a single scalar stability prediction for each protein, produced by a model that combines sequence-driven representations with global physicochemical knowledge. This multimodal approach allows LoMuS to leverage the strengths of deep language models and domain-specific features, leading to improved stability prediction performance as demonstrated in the Results section.

We obtain ESM-2 from the HuggingFace Transformers library (ensuring consistent tokenization and model weights) ^1^ [23]. Each input amino acid sequence is first tokenized with ESM-2’s tokenizer, which adds special tokens (start `<cls>` and end `<eos>`) and maps characters to token IDs. Each tokenized sequence is then fed into the ESM-2 encoder to produce a contextual embedding for each residue (plus the special tokens). Given an input amino acid sequence of length *L*, tokenization yields *L* + 2 tokens after adding `<cls>` and `<eos>`. The ESM-2 encoder (hidden size *d* = 1280) outputs hidden states *H* ∈ *ℝ*^(*L*+2)*×d*^. For downstream pooling we discard the special tokens and retain residue embeddings only, forming *X* ∈ ℝ^*L×d*^.

## Datasets

To systematically assess the proposed model architecture, we utilize different publicly available datasets from large-scale to out-of-distribution test sets. We also evaluate our model for per-protein mutational scanning usecases to support rational protein engineering. In this section, we discuss each dataset with their uniqueness and highlight our preparation steps.

### Tsuboyama Mega-scale Folding-Stability Dataset

We use the cDNA-display proteolysis dataset from Tsuboyama et al. [48]. It compiled and reported the absolute folding free energy Δ*G* for approximately 7.7 *×* 10^5^ sequences, largest publicly available protein stability dataset to the best of our knowledge, under uniform conditions near pH 7.4 and 298K. These proteins span across 331 natural and 148 *de novo* domains. Proteolysis is carried out with trypsin and chymotrypsin, and a Bayesian kinetic model estimates each sequence’s *K*_50_ and converts it to Δ*G*. Quality filters remove sequences with unreliable dynamic range or evidence of folded-state cleavage, yielding a large and internally consistent corpus of Δ*G* measurements for stability modeling [48].

We adopt a wild-type domain aware splitting protocol for training and testing of our proposed model commonly used in proteomics research following ESMtherm [18]. Contrary to the random mechanism, domain-based splitting considers keeping all mutant- and wild-type sequences into the same partition. This indicates that the entire domains are held out for testing and validation. Finally, the train, validation, and test sets contain 379, 577, 47, 414, and 100, 794 sequences, respectively. Summary statistics for different partitions are reported in the Supplementary S1.3.1. We keep Δ*G* as a continuous target without binarization. Fig. 3(A) demonstrates the stability value distributions for all splits. Overall, this dataset evaluates the generalization capacity of the model to novel domains rather than additional mutants from the same training domains.

**Fig. 3.**
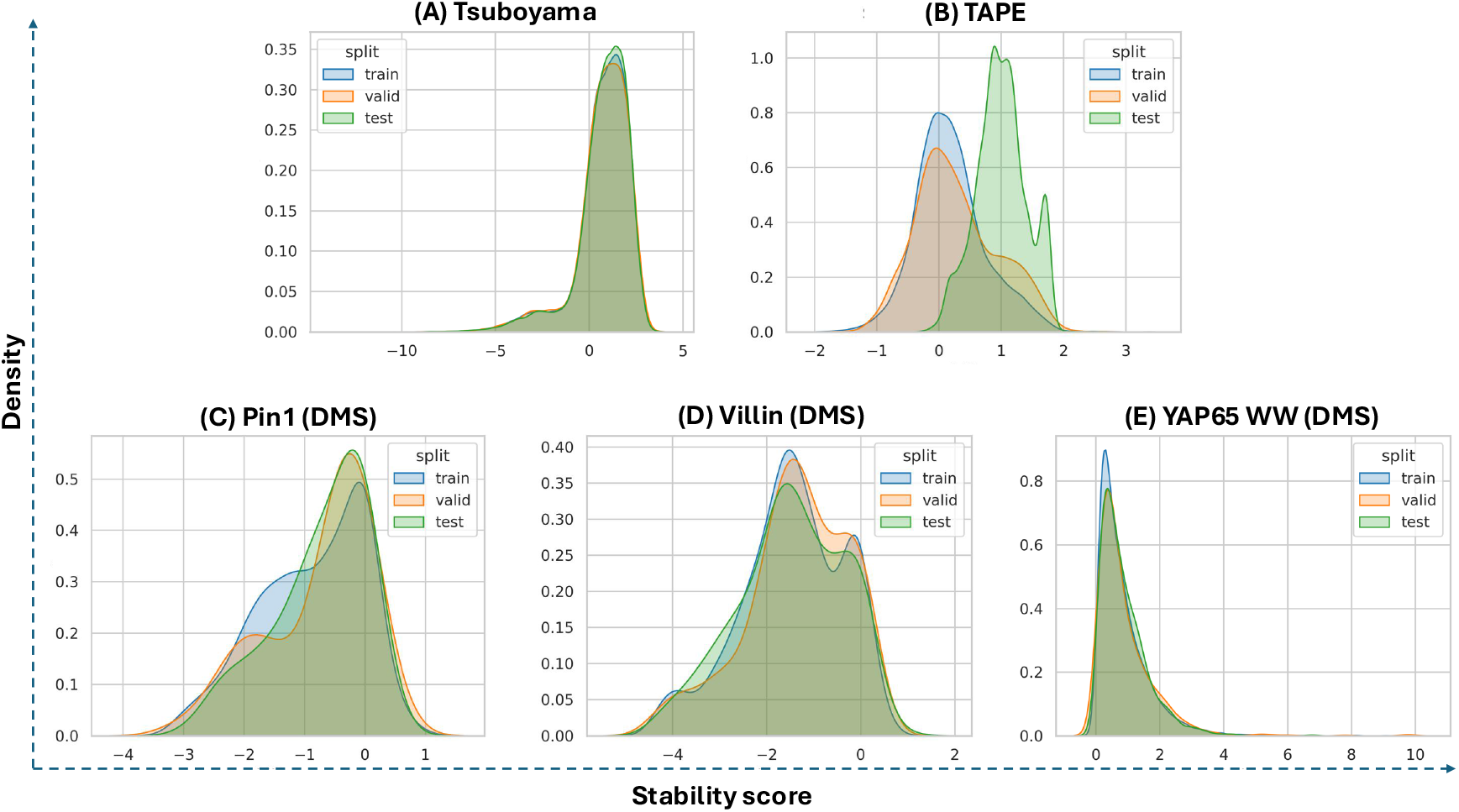
protein folding stability label distributions by train, validation and test splits across various benchmarks. (A) Tsuboyama Δ*G* mega-scale folding stability dataset [48]. (B) TAPE stability benchmark [25] exhibits a deliberate distribution shift, with the test set centered at a higher mean and lower variance than train and validation, emphasizing the challenge of generalization. (C,D and E) Per-protein deep mutational scanning (DMS) datasets for Pin1, Villin, and YAP65 WW (from left to right).

### TAPE Stability Benchmark

Next, we evaluate our model’s generalization capability when applied to the stability values situated outside the training distribution. We adopted a stability benchmark dataset from TAPE [25] where the underlying experimental measurements are obtained from the large scale protein design and stability study by Rocklin et al. [49]. The train and validation come from four rounds of experimental measurements across many candidate proteins, while the test set is built from seventeen Hamming-1 neighborhoods centered on promising proteins identified during those rounds [25]. The test examples stay near optimized scaffolds. This introduces a conscious label-distribution shift in the test set relative to the training and validation set. Fig. 3(B) shows the distribution shift in stability label statistics, where the test set shows a higher mean (1.002) and lower variance (0.409) than train (0.179 *±* 0.566) and validation (0.279 *±* 0.656) sets (Supplementary S1.3.2). Since the test sequences are close mutational neighbors of top candidates, the naive finetuning over the splits emphasizes learning local landscape around optimized parents instead of learning optimal solution, thus encourages overfitting. This design choice ensures that the optimal model must learn most salient and relevant features to molecular stability.

### Deep Mutational Scanning (DMS) (per-protein)

We include a per-protein stability benchmark evaluation drawn from three deep mutational scanning (DMS) datasets of natural proteins used in the UniRep study: the human YAP65 WW domain, the Villin headpiece, and Pin1 [6]. This benchmark provides a focused test set to compare generalizability across mutations within the context of a single protein, complementary to the broad cross-protein stability dataset as previously discussed. We also situate these tasks in the ProteinGym landscape, which aggregates hundreds of DMS assays across diverse proteins and experimental setups [50], to emphasize their relevance as representative stability DMS challenges.

Following UniRep’s protocol [6], we generate 80/10/10 train–validation–test splits for each protein, using the same fixed random seed and holding out the test set entirely until the final evaluation. Label distribution for each protein is reported in Fig. 3(C, D, E). Overall, we observe different skewness in the label distributions across proteins. This ensures a consistent comparison to the UniRep baseline and a fully independent test for each protein.

Each protein is considered as an independent task using the same pipeline. We parse the provided sequences and their experimental stability scores (numeric labels), sanitize sequences to the standard 20 amino acids, and compute the per-sequence physicochemical feature vector. We then apply z-score label normalization per protein (fit on that protein’s training split and applied to its validation and test splits) to standardize the stability scale. Summary statistics for the three protein datasets are provided in Supplementary S1.3.3.

## Results and Discussion

In this section, we analyze the efficacy of our proposed model compared with state-of-the-art benchmark approaches in three different settings. Our result shows that LoMuS’ multimodal strategy yields substantial gains in protein stability prediction across diverse benchmark datasets.

### Generalization across Held-out Domains

we evaluate the generalizability of our model in the out-of-distribution held-out protein domains. To this purpose, we utilize a domain-aware splitting protocol in the Tsuboyama [48] mega-scale folding-stability dataset. The measurements in the dataset represent short-domain Δ*G* under uniform assay conditions, and the splitting protocol confirms no overlapping proteins in the train-test splits belonging to the same domain. Our proposed model, LoMuS, outperformed the ESMtherm [18] baseline by an absolute *∼* 10% in the spearman’s rank correlation (*ρ*) metric, reported in Fig. 4(A, top-panel), specifically LoMuS achieved *ρ* of 0.75 compared to ESMtherm’s 0.65. Next, we visualize and compare the achieved performance using the parity plot containing 100, 794 test sequences. Fig. 4(B, top-panel) highlights the quality of the prediction scores and reports mean-squared error and root mean-squared error of 0.68 and 1.04, respectively. Further analysis of the errors made by the model, in Fig. 4(C, top-panel), shows that the absolute difference between the predicted vs. ground truth stability score of *>* 1.0 is occurred only for a small fraction of test samples (*≈* 22%). Overall, our proposed model sets a new baseline in the Mega-scale PROTEIN folding stability prediction benchmark.

**Fig. 4.**
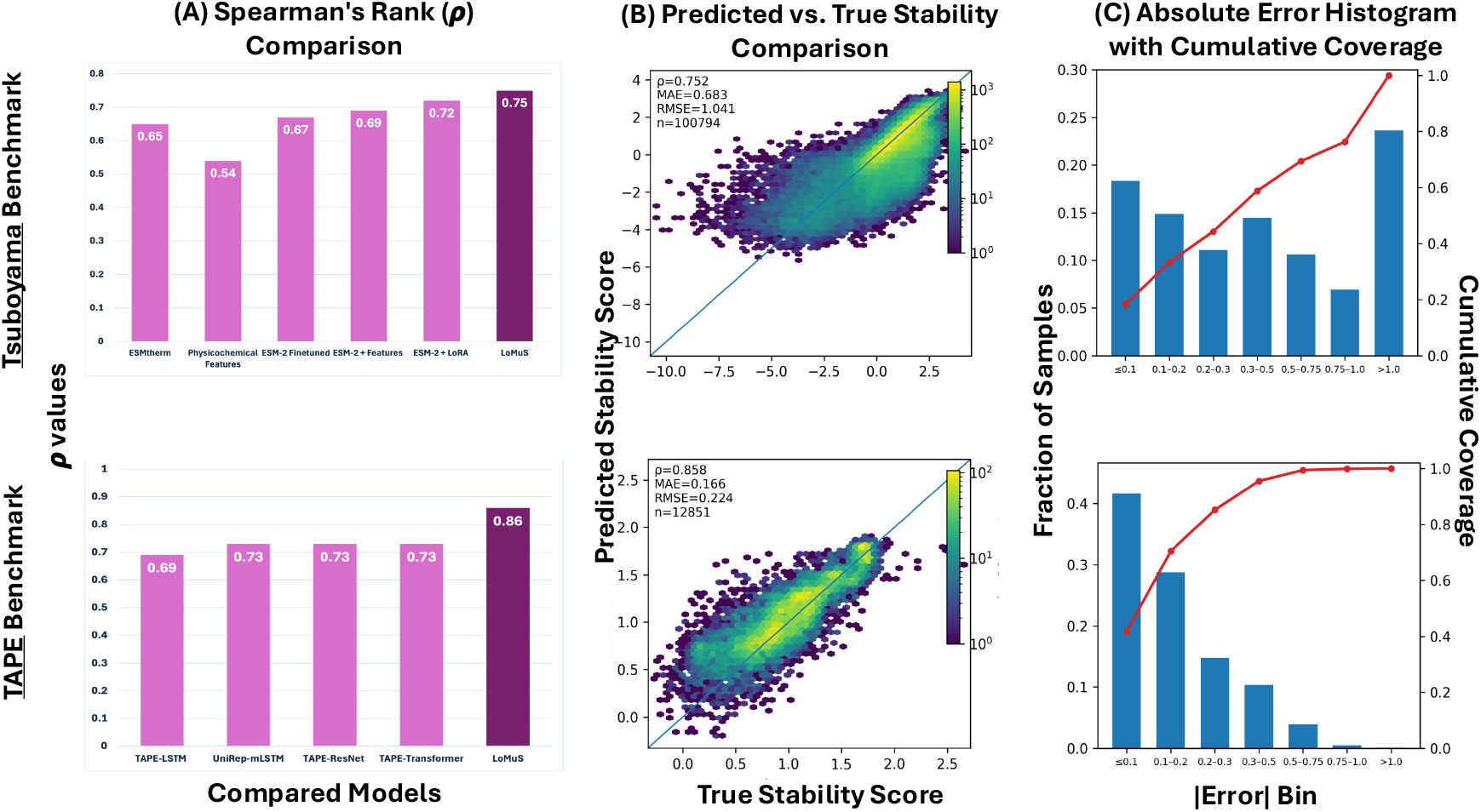
The figure highlights results across two benchmark datasets Tsuboyama [48] (top-panel) and TAPE [25] (bottom-panel). (A) Spearman’s rank correlation comparison across different models. (B) Predicted vs. ground truth stability scores comparison with spearman’s rank correlation (*ρ*), mean-squared error (MAE), root mean-squared error (RMSE) and number of test sequence (n). (C) Absolute error histogram with cumulative coverage across multiple bins. This demonstrate the superiority and quality of our proposed approach across held-out domains settings and out-of-distribution generalization.

### Generalization across Out-of-distribution Stability Labels

In our second evaluation setting, we aim to find LoMuS’s effectiveness in predicting out-of-distribution protein stability scores. We utilize TAPE [25] stability benchmark datasets and the same splits reported in the paper, ensuring a fair comparison and no-data leakage. LoMuS outperformed TAPE benchmark models by an absolute *∼* 13% in the spearman’s rank correlation metric (*ρ*), summarized in Fig. 4(A, bottom-panel). TAPE reports performance scores for models with and without self-supervised pretraining in various settings. In the original benchmark, the best pretrained Transformer and ResNet both achieved *ρ* of 0.73 on the independent test set, while the pretrained LSTM reaches 0.69. Another benchmark, UniRep (mLSTM) [6], is in the same ballpark (*ρ* = 0.73). LoMuS obtains spearman’s *ρ* of 0.86 on the TAPE stability test split, Fig. 4(A, bottom-panel), a substantial improvement over the widely cited 0.73 baseline. This closes a large portion of the remaining headroom on this task and suggests that fusing lightweight physicochemical descriptors with a LoRA adapted PLM backbone provides a complementary signal beyond sequence-only naive fine-tuning. Further analysis utilizing the parity plot and absolute-error distribution in Fig. 4(B, bottom-panel) and Fig. 4(C, bottom-panel), respectively, demonstrates the quality of the predicted stability scores across 12, 851 test sequences. In particular, less than 20% of the test samples have an absolute error greater than 0.2. Overall, LoMuS shows effectiveness and promise in out-of-distribution generalization settings, across both held-out domains and stability labels.

#### Ablation Study

To understand the robustness of our model architecture and the contributions of the multimodal representations, specifically physicochemical properties, we performed a systematic ablation study. Fig. 4(A, top-panel) compares the spearman’s rank correlation (*ρ*) for different choices of the model, such as physicochemical features only mode, naively finetuned ESM-2 backbone, and systematically incorporating LoRA with features. A features only mode that removes the PLM and uses only the physicochemical descriptors attains a spearman’s *ρ* of 0.54. As a strong sequence baseline, an ESM-2 finetuned model reaches 0.67 of *ρ*, which is close to the reported performance of ESMtherm on the same dataset. Adding the features on top of ESM-2 produces a modest gain to *ρ* = 0.69, while using only LoRA on top of ESM-2 yields a larger improvement to 0.72. The best performance is obtained when both components are used together, with ESM2+LoRA+features reaching *ρ* = 0.75, an absolute increase of about 10% over the ESMtherm baseline. We perform this ablation on the Tsuboyama Δ*G* dataset because it is the largest dataset considered in this work, which makes it more reliable for building a robust architecture. Overall, the pattern suggests that the features and LoRA provide complementary information and that the full LoMuS configuration is needed to fully exploit the multimodal representation.

We also perform the same ablation study on the TAPE stability dataset, results in Table 1. Starting from the ESM-2 finetuned baseline (*ρ* = 0.710), adding only the global physicochemical features increases performance to 0.754. Introducing LoRA gives a much larger gain, with ESM-2 + LoRA reaching *ρ* = 0.853. With the updated results, the full multimodal variant ESM-2 + features + LoRA achieves the best performance (*ρ* = 0.858), indicating that the feature branch provides an additional complementary improvement when combined with LoRA on TAPE. While this gain over ESM-2 + LoRA is modest, it is consistent with the idea that the physicochemical features can add signal beyond the sequence representation once the backbone is efficiently adapted. However, the TAPE stability benchmark has a known distribution shift between its training and test sets, and it is also much smaller than the Tsuboyama Δ*G* dataset. We therefore view the Tsuboyama ablation, where both features and LoRA contribute complementary gains on more than seven hundred thousand variants, as the primary and more reliable piece of evidence. The TAPE ablation mainly reinforces that LoRA is a key driver of performance, while the benefit of the physicochemical features can depend on dataset size and distribution.

**Table 1.** Ablation results on the TAPE stability dataset.

| Metric | ESM-2<br>finetuned | ESM-2 +<br>features | ESM-2 +<br>LoRA | LoMuS |
| --- | --- | --- | --- | --- |
| Spearman’s $\rho$ | 0.710 | 0.754 | 0.853 | **0.858** |

### Per-protein Evaluation for Protein Screening

Per-protein engineering is a critical component in therapeutics and drug design. To address this challenge, we next attempt to understand the effectiveness of our model across different individual protein settings. We utilize deep mutational scanning (DMS) dataset for three naturally occurring proteins. We compare our results with those of UniRep [6], a sequence-only multiplicative LSTM (mLSTM) pretrained with large-scale self-supervision, which established a stability benchmark in DMS across multiple proteins.

LoMuS achieves a spearman’s rank correlation (*ρ*) of 0.89, 0.97 and 0.78 in the three test sets for Pin1, Villin and YAP65 WW, respectively. Our model outperforms UniRep or performs on par, notably obtaining an absolute *∼* 11% performance gain for Villin (Table 2). Fig. 5 visualizes our model’s prediction quality using the parity plot (top-panel) and reports absolute prediction errors with cumulative coverage (bottom-panel) compared to the stability values of the ground-truth in the test sets across these three proteins. In all protein settings, less than 5% of the samples have an absolute error difference greater than 1.0.

**Table 2.** DMS per-protein results on the test set (Spearman’s *ρ*).

| Protein | UniRep (mLSTM) | LoMuS |
| --- | --- | --- |
| YAP65 WW | **0.78** | 0.776 |
| Villin | 0.86 | **0.966** |
| Pin1 | 0.89 | **0.894** |

**Fig. 5.**
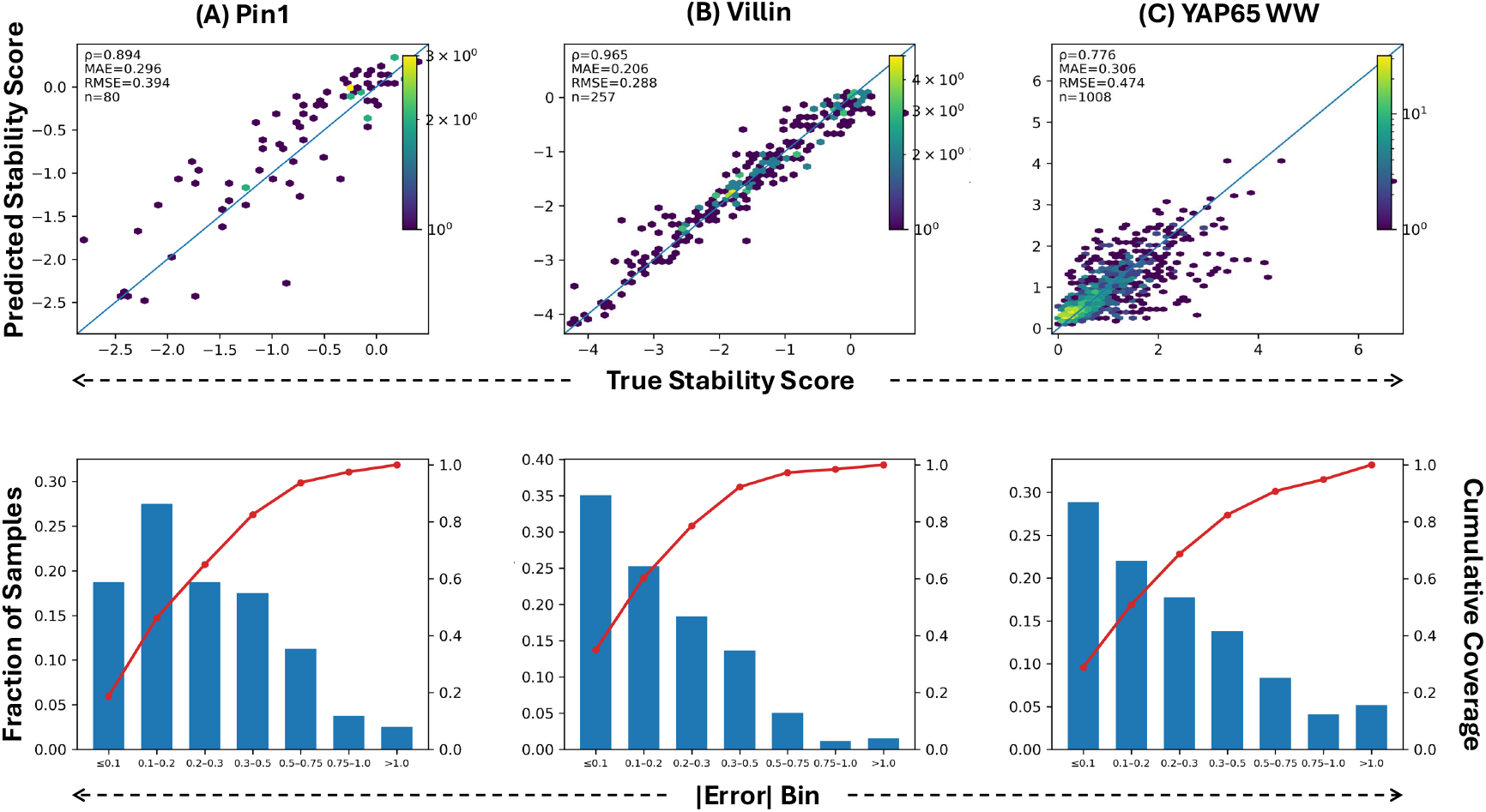
Model performance per-protein deep mutational scanning analysis for (A) Pin1, (B) Villin, and (C) YAP65 WW proteins. This demonstrate the overall quality of the model prediction scores (top panel) and low error difference (bottom-pane) across specific proteins.

Overall, our fused features captured stability determinants that a sequence-only model missed. Even when the baseline was already high, as in the case of Villin, incorporating global descriptors maintained or slightly improved performance without reducing generalization.

## Conclusion and Future Work

Designing stable molecules is a fundamental challenge in protein engineering and therapeutics. In this work, we propose a multimodal deep learning framework, LoMuS, to predict protein stability scores given the primary amino acid sequence as input. We conducted a rigorous evaluation across three different settings that demonstrate the generalizability of LoMuS in various out-of-distribution datasets. The core of the LoMuS’s architecture leverages a state-of-the-art PLM. It implicitly benefits the model with learned evolutionary signals and structural patterns from a large corpus of protein sequences. At the same time, the inclusion of known physicochemical descriptors provide orthogonal information compared to the embeddings. Finally, the low-rank adaptation with the careful design of a multimodal fusion network enables LoMuS to generalize to novel domains and to label shifts. A reliable in silico stability predictor like LoMuS can facilitate rational design cycles in protein engineering workflows. We also anticipate that the principles of LoMuS, which combine pretrained sequence embeddings with carefully selected descriptive features in a low-rank fine-tuning framework, can be extended to other protein modeling tasks where interpretability and data efficiency are important.

With the convincing success of our proposed model, we aim to extend the multimodal formulation to richer input modalities when available. A unified stability predictor by integrating lightweight structural summaries, wet-lab or computationally derived, contact patterns, solvent exposure, and/or evolutionary context might enrich model’s confidence and improve performance. Furthermore, the physicochemical indices used in LoMuS were calibrated on folded proteins; their relevance to disordered proteins is uncertain. In practice, intrinsically disordered proteins (IDPs) or regions fed into LoMuS might be predicted as “unstable” due to high instability index or low hydrophobicity, however, this does not capture the biology of disorder. Therefore, Expanding and validating the model on a wider range of protein types such as membrane proteins, multi-domain assemblies, and intrinsically disordered proteins will be another future step to consider.

In another direction, we aim to advance our proposed model to real world protein engineering practice. Instead of focusing only on stability regression, future work could engage the same multimodal recipe on other tasks such as solubility, expression levels, binding affinity or enzyme activity, as well as on multi task setups where several properties are learned and predicted jointly. In summary, LoMuS advances the field towards predictive models of protein stability that are both highly accurate and explainable, thus supporting rational protein engineering and our understanding of sequence–stability relationships.

## Computational requirements

Experiments are performed on a single NVIDIA A100 GPU on the GAIVI cluster, with 8 CPU cores and 32 GB RAM. Training is also feasible on a 12 GB NVIDIA GPU when using LoRA, mixed precision, and gradient accumulation. We used Python 3.10, PyTorch 2.*x* with CUDA 11.8 or newer, Hugging Face Transformers 4.40 or newer, PEFT for LoRA, SciPy, scikit-learn, NumPy, pandas, and Biopython for basis descriptors.

## Competing interests

No competing interest is declared.

## Author contributions statement

S.I. conceived and conducted the experiments, analyzed the data, and wrote the initial draft of the manuscript. A.S. provided additional assistance with experiments and analysis. A.K supervises the scientific work by formal analyzing the results, reviewing, and editing the drafts. All authors have reviewed the final manuscript.

## Acknowledgments

This work was supported by the faculty seed funding from the University of South Florida, FL, USA. We acknowledge the GAIVI computing cluster at USF for providing access to A100 GPUs via SLURM, which enabled all training and evaluation runs.

## Supplementary Information

## Abbreviations used in this paper

ML: Machine Learning
DL: Deep Learning
PLM: Protein Language Model
MLPs: Multi-layer Perceptrons

## Six global physicochemical features

- Sequence length: Number of standard amino acids in the protein.
- Molecular weight: Total mass of the protein in Daltons, computed from the sequence composition.
- Isoelectric point (pI): The pH at which the protein has net zero charge. We estimated from amino acid pKa values.
- GRAVY (Grand Average of Hydropathy): The mean hydropathy value of the sequence, indicating overall hydrophobicity or hydrophilicity.
- Aliphatic index: A measure of the volume of aliphatic side chains (Ala, Val, Ile, Leu) in the protein, which is empirically correlated with thermostability.
- Instability index: An empirical measure of in vitro stability, where a value <40 predicts a stable protein and >40 predicts an unstable protein.

### Reproducibility

All experiments are conducted with reproducibility in mind. We use a fixed random seed for weight initialization and data shuffling, ensuring that results are deterministic. Additionally, we preserve the original order of sequences and labels as provided. For example, if sequences are listed in a FASTA file with a corresponding label list, we do not randomize their pairing. Each input sequence is always matched to the correct label by index. This deterministic ordering (no inadvertent shuffling misalignment) guarantees that the same sequence-label mapping is used across runs, and that cross-validation splits are consistent. By aligning sequences and labels deterministically and fixing the random seed, we facilitate exact reproducibility of model training and evaluation.

## Summary of Dataset Statistics

We summarize key descriptive statistics for all datasets used in this study to provide context on dataset size and label distribution characteristics across splits. For each benchmark, we report the number of sequences, mean and standard deviation of the stability labels, highlighting differences in scale, distribution shifts, and sequence constraints across datasets.

### Tsuboyama Mega-scale Folding-Stability Dataset [48]

**Table S1.** Summary statistics for *Tsuboyama* dataset splits.

| Split | Number of Samples | Average | SD (±) |
| --- | --- | --- | --- |
| Train | 379,577 | 0.722 | 1.483 |
| Validation | 47,414 | 0.707 | 1.487 |
| Test | 100,794 | 0.739 | 1.453 |

### TAPE Stability Benchmark[25]

**Table S2.** Summary statistics for TAPE Stability dataset splits.

| Split | Number of Samples | Average | SD (±) |
| --- | --- | --- | --- |
| Train | 53,614 | 0.179 | 0.566 |
| Validation | 2,512 | 0.279 | 0.656 |
| Test | 12,851 | 1.002 | 0.409 |

### Deep Mutational Scanning (DMS) (per-protein) [6]

**Table S3.** Summary statistics for DMS proteins datasets splits.

| Protein | Split | Number of Samples | Average | SD (±) |
| --- | --- | --- | --- | --- |
| Pin1 | Train | 642 | 0.876 | 0.871 |
|  | Validation | 80 | 0.756 | 0.864 |
|  | Test | 80 | 0.729 | 0.770 |
| Villin | Train | 2,054 | 1.467 | 1.096 |
|  | Validation | 257 | 1.368 | 1.090 |
|  | Test | 257 | 1.486 | 1.114 |
| YAP65 | Train | 8,059 | 0.855 | 0.807 |
|  | Validation | 1,008 | 0.897 | 0.925 |
|  | Test | 1,008 | 0.866 | 0.723 |

## Footnotes

1 https://huggingface.co/facebook/esm2_t33_650M_UR50D

## Notes

### Competing Interest Statement

The authors have declared no competing interest.

